# Epistatic Modifiers Influence the Expression of Continual Flowering in Strawberry

**DOI:** 10.1101/2022.03.15.484381

**Authors:** Helen Maria Cockerton, Charlotte Florence Nellist, Timo Hytönen, Suzanne Litthauer, Katie Hopson, Adam Whitehouse, Maria Sobczyk, Richard Jonathan Harrison

## Abstract

Previous work within the community led to the identification of a single dominant allele that controls the everbearing trait. However, frequent observations have indicated that crosses do not segregate in a Mendelian fashion, as would be expected for a trait controlled by a single dominant gene. Therefore, it was hypothesised that one or more unidentified epistatic alleles interact with the major gene.
A GWAS was conducted on 587 June-bearers and 207 everbearers to assess the genetic components associated with flowering habit. The segregation ratios of parental strawberry lines with known phenotypes were used to validate the identified alleles.
Three loci including the known, major *FaPFRU* locus and two epistatic modifiers were identified. These modifiers function as enhancers of the everbearing trait in individuals containing a single copy of the *FaPFRU* everbearing allele and appear to be functionally redundant. Principally, heterozygous individuals required the presence of two modifying alleles in order to allow expression of the everbearing trait.
Inclusion of the epistatic alleles improved the prediction of everbearing segregation ratios, beyond that of a single allele model, however, a large proportion of the variation remained unexplained. Future work should identify the additional repressor and enhancer modifiers not identified here. Discovering the genetic components controlling the everbearing trait can enable genetic informed strawberry improvement.

**Society Impact Statement:** Until the 1970s, the majority of commercial strawberry varieties produced a single bloom of flowers. However, continuously flowering, everbearing strawberries are now routinely cultivated and use is increasing. Indeed, introgression of the everbearing flowering trait can lead to economic benefits for growers through the production of a continual crop from the same plant. Utilising genetic guided improvement has the power to streamline everbearing generation. As such, utilising the genetic markers reported here, can help to identify everbearing individuals at an early time point in the breeding process. Furthermore, these markers can help to improve the predictions of progeny segregation ratios.

## Introduction

Plants that grow in constant environments are able to achieve high reproductive success through continuous flower production. However, angiosperms that grow in temperate environments must flower within a restricted time period; flower too early and risk damage due to late frosts, flower too late and seed cannot be developed and dispersed before the onset of winter (Gaudinier & Blackman, 2020). To maximise the chance of reproductive success, a plant must also synchronise its life cycle with that of mutualistic organisms; both pollinators and dispersers (Krizek & Fletcher, 2005). As such, plants have evolved to modify the timing of flower emergence in response to different environments. Plants that grow in temperate environments need to tightly regulate flower production in order to achieve successful reproduction within a limited time frame. In order to regulate flowering time, temperate plants have evolved the ability to measure the season. Thus temperate adapted plants use environmental cues that cycle reliably throughout the year; via the length of the day (photoperiod) and through a prolonged period of chilling where vernalisation generates a “winter memory” (Andrés & Coupland, 2012; Bouché *et al.*, 2017). Indeed, expansion of plants from tropical to temperate regions required the ability to control flowering time and as such “winter memory” has co-evolved numerous times, independently, across multiple angiosperm families (Bouché *et al.*, 2017). By contrast, non-seasonal climates do not have annual fluctuations and thus there is no selection pressure on angiosperms to flower within a specific time frame.

Plants in the *Fragaria* genus are widely distributed across many different geographic locations and environments (Darrow, 1966; Liston *et al.*, 2014). Seasonally flowering strawberry plants, identified as short-day or June-bearers, initiate flower buds in autumn and the flowers emerge in the following spring. Flower induction requires short day conditions, when the photoperiod is less than about 14 hours in length. However, temperature can modulate the short day flowering response whereby strawberry plants are induced to flower in long photoperiods at cool temperatures (<15 °C) (Guttridge, 1985; Mookerjee *et al.*, 2013; Rantanen *et al.*, 2015). By contrast, strawberry genotypes that produce continuous or repeated flowers are termed “everbearers”. Temperature and light periodicity also influence flowering time in everbearing strawberries, but after flower induction, everbearing strawberry plants will continuously produce flowers in both short and long days (Bradford *et al.*, 2010; Koskela *et al.*, 2012). This continual flowering phenotype is of particular interest for strawberry breeders and growers due to the potential benefit of increased yield from a single planting for the duration of the growing season.

Everbearing is a relatively new trait in strawberry cultivation, and the majority of historical commercial production has focused on June-bearers. The first documented introgression of the everbearing trait was from a wild *Fragaria virginiana* ssp. *glauca* from the Wasatch Mountains in 1955 (Bringhurst & Voth, 1980). It may be hypothesised that colder mountain conditions (<15 °C) induced flowering, which removed the selection pressure to maintain the short day flowering mechanism and thus everbearing plants evolved. Introgression of the everbearing trait into commercial strawberry cultivars has allowed continual fruit production to be extended across the season. The Wasatch source of everbearing has been used internationally and is thought to represent the main genetic source of everbearing in temperate cultivated strawberries (Simpson, 1993; Faedi *et al.*, 2002; Zurawicz & Masny, 2002; Salinas *et al.*, 2017). Before the commercialisation of everbearing strawberry varieties and the associated improvement of fruit quality, it had been noted that significant refinement of everbearing genetics was required (Bringhurst & Voth, 1984).

A single major dominant locus *FaPFRU* was found to control both everbearing and runnering (clone plants produced from horizontal stems) in octoploid strawberries. Individuals containing the dominant allele(s) of this locus produced up to 20 times more inflorescences and up to 16 times fewer primary runners (Gaston *et al.*, 2013). The *FaPFRU* locus has since been fine mapped and two copies of the identified recessive haploblock were required to produce a short day variety (Verma *et al.*, 2017b). Furthermore, a SSR (simple-sequence repeat) marker has been validated to colocalise with the *FaPFRU* locus across wider germplasm (Salinas *et al.*, 2017). This marker has allowed the use of marker assisted breeding to incorporate everbearing into high fruit quality lines without incurring phenotyping costs. Identifying a functional marker in the causative gene or allele would allow for highly accurate selection of the trait and negate the requirement for maintenance of non-desirable individuals. However, it has since been discovered that the “single dominant gene” controlling the everbearing trait does not segregate in a straightforward Mendelian fashion in all biparental crosses (Lewers *et al.*, 2019). Indeed, it was hypothesised that the allele might be controlled by at least two epistatic alleles that interact with *FaPFRU* to modify the expression of the everbearing trait (Lewers *et al.*, 2019). Here we provide evidence to confirm their original hypothesis and identify two new epistatic loci that are involved in the expression of continual flowering in strawberries.

## Materials and Methods

### Plant Material

The in-house NIAB EMR strawberry genomic database was mined to study the genetic control of the everbearing trait using a Genome Wide Association Study (GWAS). A total of 794 strawberry breeding lines and cultivars had genotypic data available with a known flowering habit status. The population represents modern released cultivars (post 1980’s) and NIAB EMR advanced breeding lines generated between 2000 and 2018.

### Phenotyping

Flowering habit was scored as a binary trait, genotypes were denoted as June-bearers or everbearers. Determination of flowering habit for each genotype was achieved through either literature searches, cultivar release information or determined based on the in-house phenotype databases. The initial in-house characterisation consisted of observing flower production in field conditions or protected substrate culture over two consecutive growing seasons (April-Oct) at NIAB EMR (51° 17′ 24.202″ N 0° 26′50.918″ E). The presence of open inflorescences and fruiting was scored on a weekly basis over the season to confirm the June-bearing or everbearing nature of each of the genotypes.

### Genotyping

Genotyping data used for the GWAS analysis was generated as part of a previous project (Nellist *et al*. 2019). The parents of the validation germplasm were genotyped as part of this work; newly unfolded strawberry leaf material was sampled for DNA extraction using the QIAamp 96 DNA QIAcube HT Kit, with buffer volumes modified for isolation of total DNA from plant tissue, as per manufacturer's instructions. DNA quality was confirmed using a spectrophotometer (NanoDrop^TM^ 2000) and DNA quantity was confirmed through Qubit analysis. All genotyping was performed on the Istraw 90k or the Istraw 35k axiom array (Bassil *et al.*, 2015; Verma *et al.*, 2017a).

### Genetic Analysis

Genome wide association analysis was conducted in Plink (Purcell *et al.*, 2007) as detailed on github https://github.com/harrisonlab/popgen/blob/master/snp/gwas_quantitative_pipeline.md. A total of 94,748 informative SNPs (single nucleotide polymorphism) were used for the analysis presented here. SNPs were filtered to include only those with a minor allele frequency of greater than 0.05, and no greater than 20 % missing values across the genotypes studied. Individuals with high heterozygosity were removed. The analysis was adjusted for population stratification through logistic regression using multidimensional scaling values as covariates.

Epistatic interactions between the QTL (quantitative trait loci) were quantified through a binomial generalised linear model, with the formula: Flowering habit ~ A + B + C + A*B + B*C. Where letters (A, B & C) denote the three genetic loci. Data from 771 genotypes were used to assess epistatic interactions as 23 individuals had “no call” for one of the three alleles.

Genotypic relationships were calculated through hierarchical cluster analysis of the Euclidean distance matrix, this data was plotted as a dendrogram using the R packages dendextend & ggplot (Wickham, 2009; Galili, 2015).

### Determining the Origin of SNPs

The origin of the SNPs of interest were investigated by assessing the presence/absence in wild octoploids using an existing dataset (Hardigan *et al.*, 2018). A dendrogram was generated by hierarchical clustering for a subset of 266 wild accessions using 16,394 markers.

### Validation of the Model using Bi-parental Crosses

Progeny from 156 crosses were screened for flowering habits; crosses had between 30 and 400 progeny. A linear model was generated between the observed and predicted percentage of everbearing individuals and values were weighted by the total number of progeny in the population. The predicted percentage of everbearing individuals was based on the genotype of the parents for one of three models: (1) the single allele model for the major allele (A), (2) the simple three allele model where one modifier (B or C) is required for heterozygous individuals for the major allele (Aa) to exhibit the everbearing trait, (3) the complex three allele model where heterozygous individuals for the major allele (Aa) require at least two of the modifiers (either BB, CC or BC) to exhibit the everbearing trait.

## Results

### The Three Gene Model for Everbearing Trait

A total of 587 June-bearing and 207 everbearing strawberry varieties were analysed in a GWAS. Three highly significant loci were detected (Figure 1a). One large effect locus was identified on chromosome 4A with the most significant marker Affx-88857614 located at 29,532,771 bp in *F. vesca* genome v4.0 (Edger et al., 2018). Hardigan et al. (2018) also identified this locus in different germplasm alongside a marker about 3.5 Mb downstream of our focal marker on the chromosome 4A, denoted hereafter as A’, this region is known to the community as the *FaPFRU* locus (Gaston *et al*., 2013). In our GWAS, A’ showed a much lower significance level than A. We identified an additional highly significant marker X at the distal end of chromosome 4A near to A’. X delineated a 2.85 Mb marker gap between A and X (Figure 1b). Significant pairwise interaction was observed between A and A’ (*z*= −2.44; *p*= 0.015) and between A and X (*z*= −5.129; *p*= 3.7 × 10^7^).

**Figure 1.**
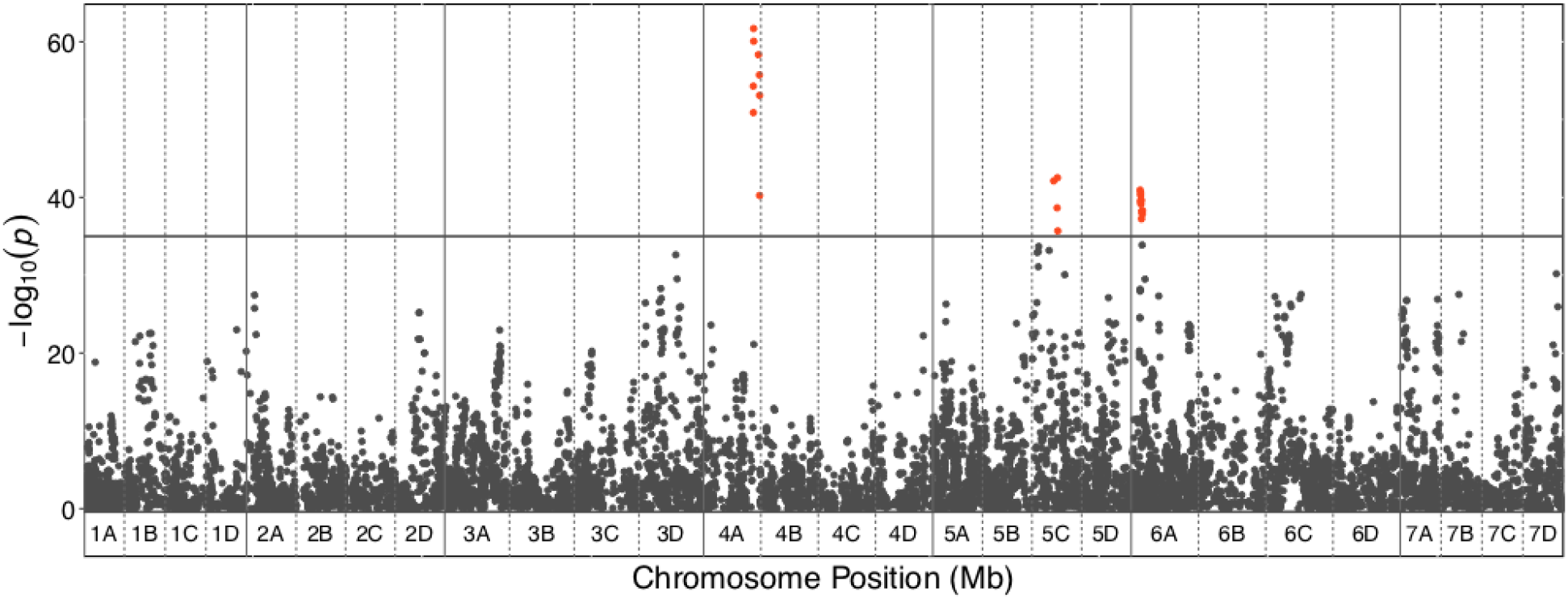
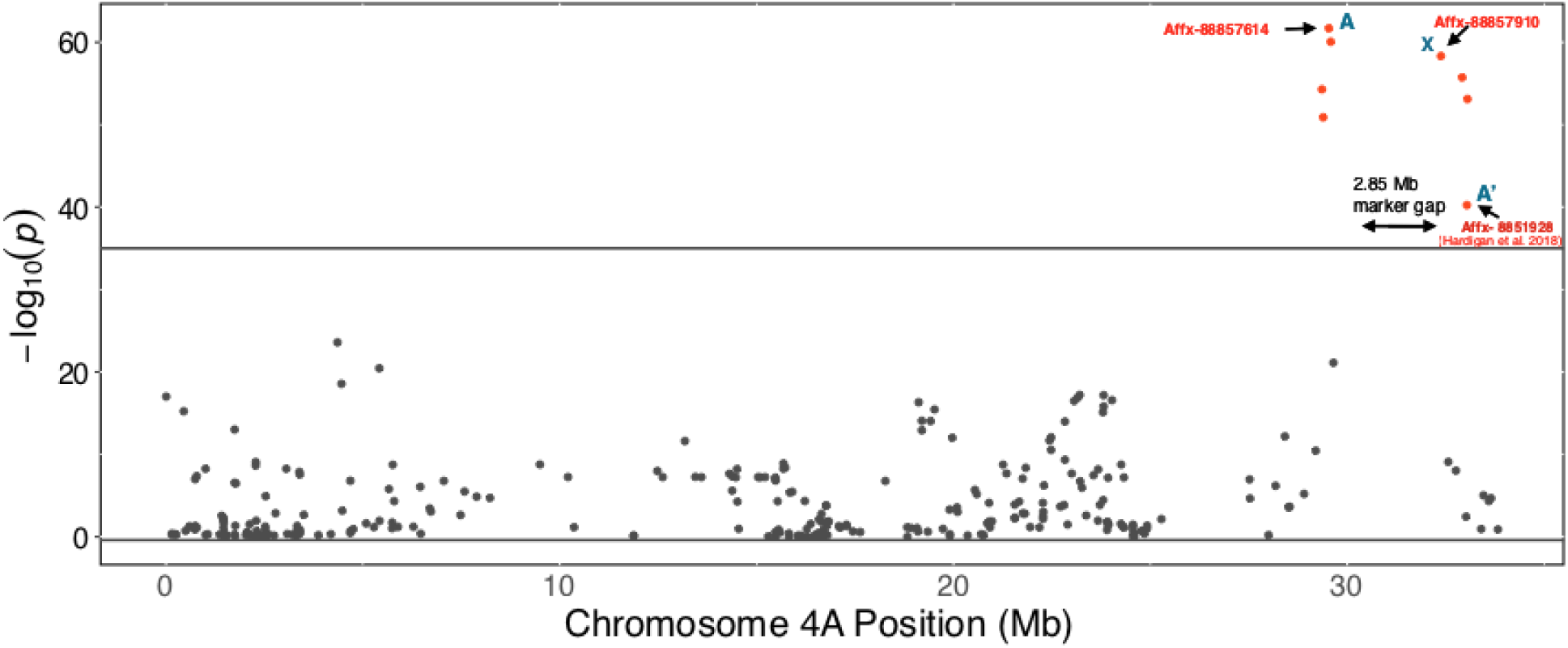
Manhattan plots showing the association of SNPs in the 794 accessions with flowering habit in octoploid strawberry. Marker positions are scaled to the *F. vesca* version 4 genome (Edger et al., 2018). Red points represent markers passing the significance threshold. Grey points represent markers beneath the significance threshold. **a** Three significant quantitative trait loci (QTL) regions identified after genome wide association analysis. Chromosomes are denoted by 1A - 7D. **b** Regional plot, zoomed in on Chromosome 4A, depicting the most significant markers, denoted as A and X and the previously identified marker by Hardigan et al. (2018), denoted as A’.

The majority of individuals (97.8%) which contained two recessive copies of the major locus (here denoted as ‘aa’) were June-bearers. The 17 everbearers that were ‘aa’ included the named cultivars ‘Diamante’, ‘Elan’, ‘Portola’ and ‘Sierra’. All individuals containing two dominant versions of the major locus ‘AA’ were everbearers (*n* = 59). Of individuals containing a single copy of the major locus ‘Aa’, 64.7 % were everbearers (*n* = 130) and 35.3 % June bearers (*n* = 71). The whole distal portion of chromosome 4A was associated with the everbearing trait, where individuals containing both dominant upper (A) and lower portions (X) gave rise to the greatest proportion of everbearing individuals (Supp. Figure 1).

Two additional loci were identified on linkage group 5C (‘B/b’; Affx-88866698 at 15,100,245 bp) and 6A (‘C/c’; Affx-88877135 at 5,285,949 bp). A significant epistatic interaction between the major locus A and the additional loci B (*p* = 0.025) and C (*p* = 0.001) was observed, but no additive interaction was found for these components (Table 1, Figure 2). The phenotypes of heterozygous genotypes for the major loci (‘Aa’) appear to be influenced by the presence of the two additional loci indicating that they function as modifiers of *FaPFRU*. A single copy of B or C is sufficient to permit the everbearing genotype to occur in some individuals. However, the majority of individuals (87 %) that did not contain one of the two modifying QTL (with the genotype ‘Aabbcc’) were June-bearers. Also ‘AaBbcc’ and ‘AabbCc’ genotypes appear to divide into two flowering types with 43.3 and 50 % of individuals exhibiting an everbearing flowering habit, respectively. There were no individuals with an ‘AAbbcc’ genotype, as such it is not clear whether ‘AA’ alone is sufficient to produce an ever-bearing phenotype. Loci B does not neighbour any candidate genes known to play a role in the flowering response, by contrast loci C is located 100 kb away from a gene encoding the photoreceptor Phytochrome A.

**Table 1.** Generalized linear model parameters for the effect of A, B and C alleles on flowering habit and pairwise interactions.

|  | Estimate | Std. Error | z value | P value | Significance |
| --- | --- | --- | --- | --- | --- |
| (Intercept) | 73.8 | 15.0 | 4.9 | $9.4 \times 10^{-7}$ | *** |
| A | -17.2 | 3.2 | -5.3 | $9.4 \times 10^{-8}$ | *** |
| B | -7.7 | 2.4 | -3.2 | 0.002 | ** |
| **C** | -12.1 | 4.0 | -3.0 | 0.003 | ** |
| **A\*B** | 1.2 | 0.5 | 2.2 | 0.025 | * |
| **A\*C** | 2.7 | 0.8 | 3.3 | 0.001 | ** |
| **B\*C** | 0.5 | 0.6 | 0.9 | 0.362 | NS |

Significance codes: 0 ‘\*\*\*’, 0.001 ‘\*\*’, 0.01 ‘\*’, NS non significance.

**Figure 2.**
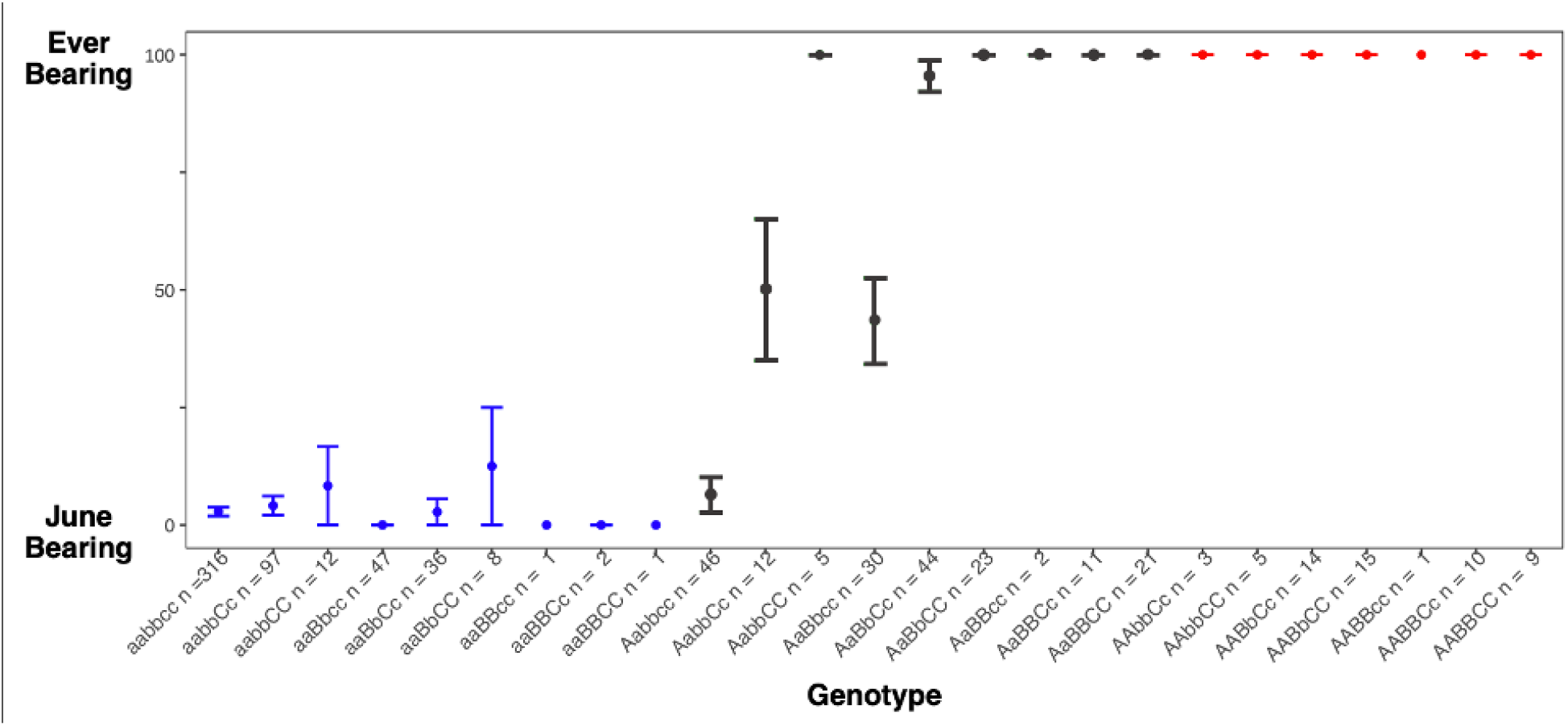
Three alleles interact to determine the flowering habit phenotype of individuals Flowering habit of each genotype represented in the population. Points represent the proportion of individuals exhibiting a June-bearing (0) or everbearing (1) phenotype. Error bars are standard errors. Loci resulting in the everbearing phenotype denoted with a capital letter (A/B/C), loci resulting in the June-bearing phenotype denoted in a lower case letter (a/b/c). Blue points represent homozygous recessive individuals for the major allele - aa. Grey points represent heterozygous individuals for the major allele - Aa. Red points represent homozygous dominant individuals for the major allele - AA. A total of 771 individuals are represented in this plot.

### Phylogeny of the Everbearing Trait

We analysed the genetic relationships between individuals within the GWAS population by generating a dendrogram (Figure 3). The everbearing trait is mainly present in one of the three clades, indicating that many of the everbearing individuals appear to be highly related. This may be due to introgression from a single source, i.e. the Wasatch source that was introduced to the University of California breeding program and later widely used across the world (Bringhurst & Voth, 1980). Some individuals in this clade appear to have “reverted” to the June-bearing phenotype. However, everbearing individuals were also found across the length of the dendrogram in the other clades.

**Figure 3.**
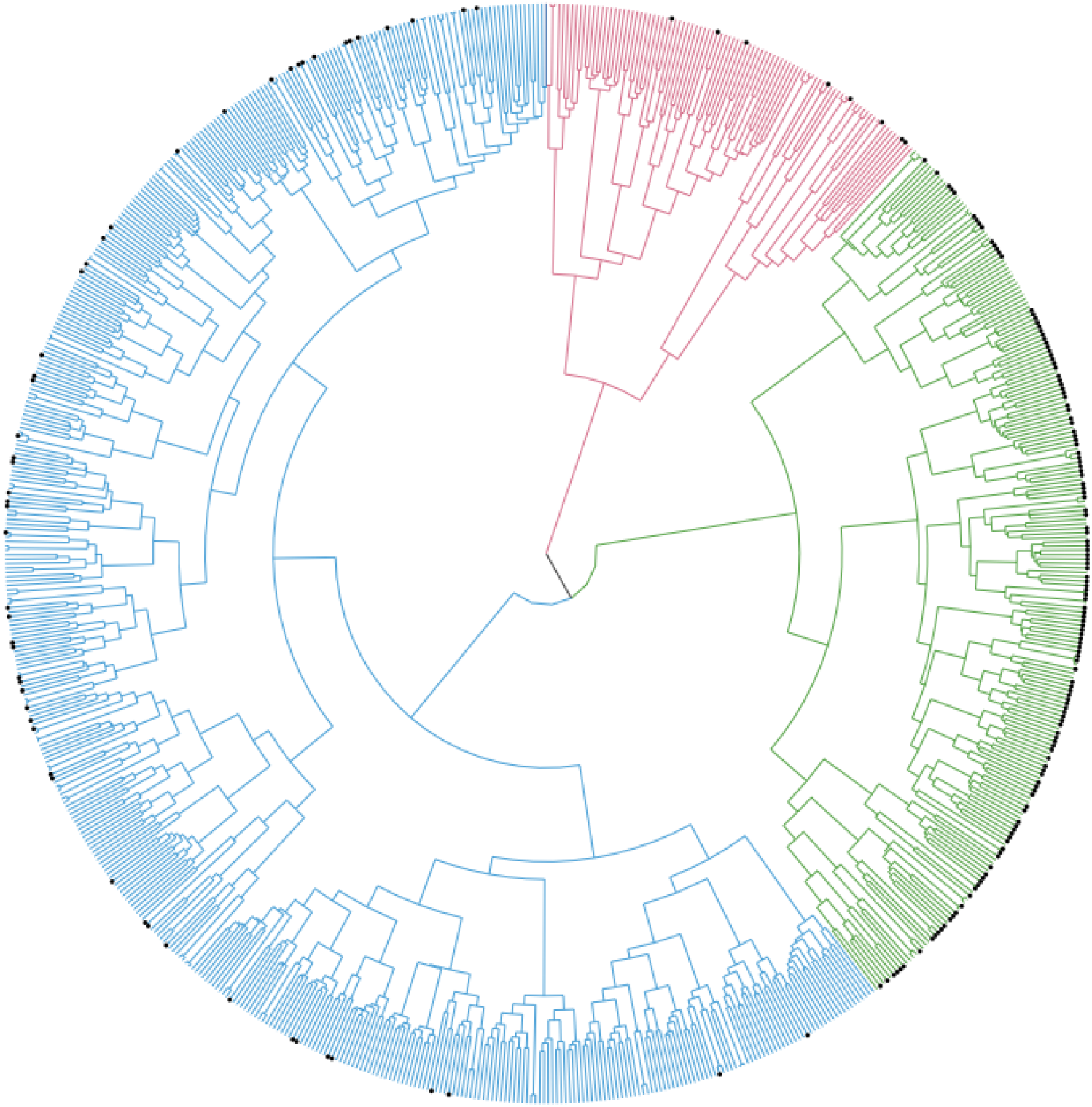
Everbearing individuals identified across all three clades of strawberry, the majority of everbearing individuals belong to a single clade. Dendrogram depicting the genetic relationship between everbearing (black points) and June-bearing (no points) cultivars. Branch colours denote the three main clusters of genotypes based on the genotypic relationship across individuals.

### Validation Crosses

We generated 156 validation crosses and compared observed versus predicted segregation ratios using three different genetic models for the everbearing trait (Figure 4). The complex three allele model (where heterozygous individuals for the major allele (Aa) require at least two of the modifiers, either BB, CC or BC, to be everbearing) has the best fit with the highest adjusted *R^2^* value of 0.33 indicating that the complex ABC model may provide a better predictive segregation ratio than the single allele model (where a single dominant A allele is sufficient) or the simple ABC model (where heterozygous Aa requires either dominant B or C). Furthermore, the equation of the complex model regression line is *y*= −8.68 + 1.01*x* which is closer (relatively) to the perfect prediction model (*y*=1*x*) than the equations of other models tested (Figure 4). Nonetheless the model does not explain all the variation in segregation ratios and it is clear there may be more unidentified alleles or other factors influencing expression of the trait.

**Figure 4.**
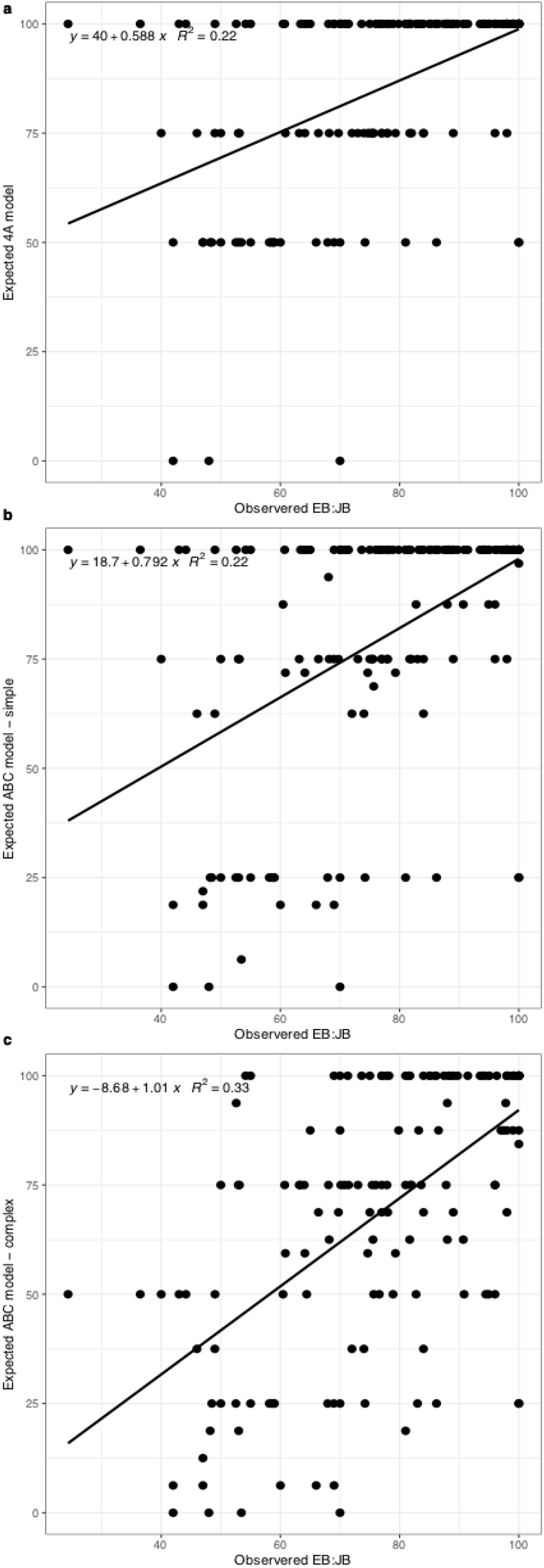
The complex three allele (ABC) model explains the most variation. Observed versus expected everbearing (EB): June-bearing (JB) segregation ratios in 156 validation crosses based on **a** the single allele model (4A), **b** the simple ABC model and **c** the complex ABC model.

### Potential Origin of the Alleles

All four alleles identified in this study were found in *F. virginiana* ssp. *virginiana*, whereas only three of the four alleles were identified in *F. virginiana* ssp. *glauca*, the original source of everbearing trait from the Wasatch mountains (Bringhurst & Voth, 1980), and in *Fragaria chiloensis* ssp. *pacifica* (Table 2). Analysis of the origin of the A, A’, B and C alleles using the data from Hardigan *et al.* (2018) revealed that the alleles are restricted to and segregating within *F. chiloensis* ssp. *pacifica*, *F. virginiana* ssp. *glauca* and *F. virginiana* ssp. *virginiana*, despite there being multiple other subspecies and populations of both *F. virginiana* and *F. chiloensis* in the dataset (Table 2). Phylogenetic analysis clearly shows that the everbearing trait stems from two discrete pools of individuals, represented by *F. chiloensis* spp. *pacifica* and *F. virginiana* (Figure 5).

**Table 2.**
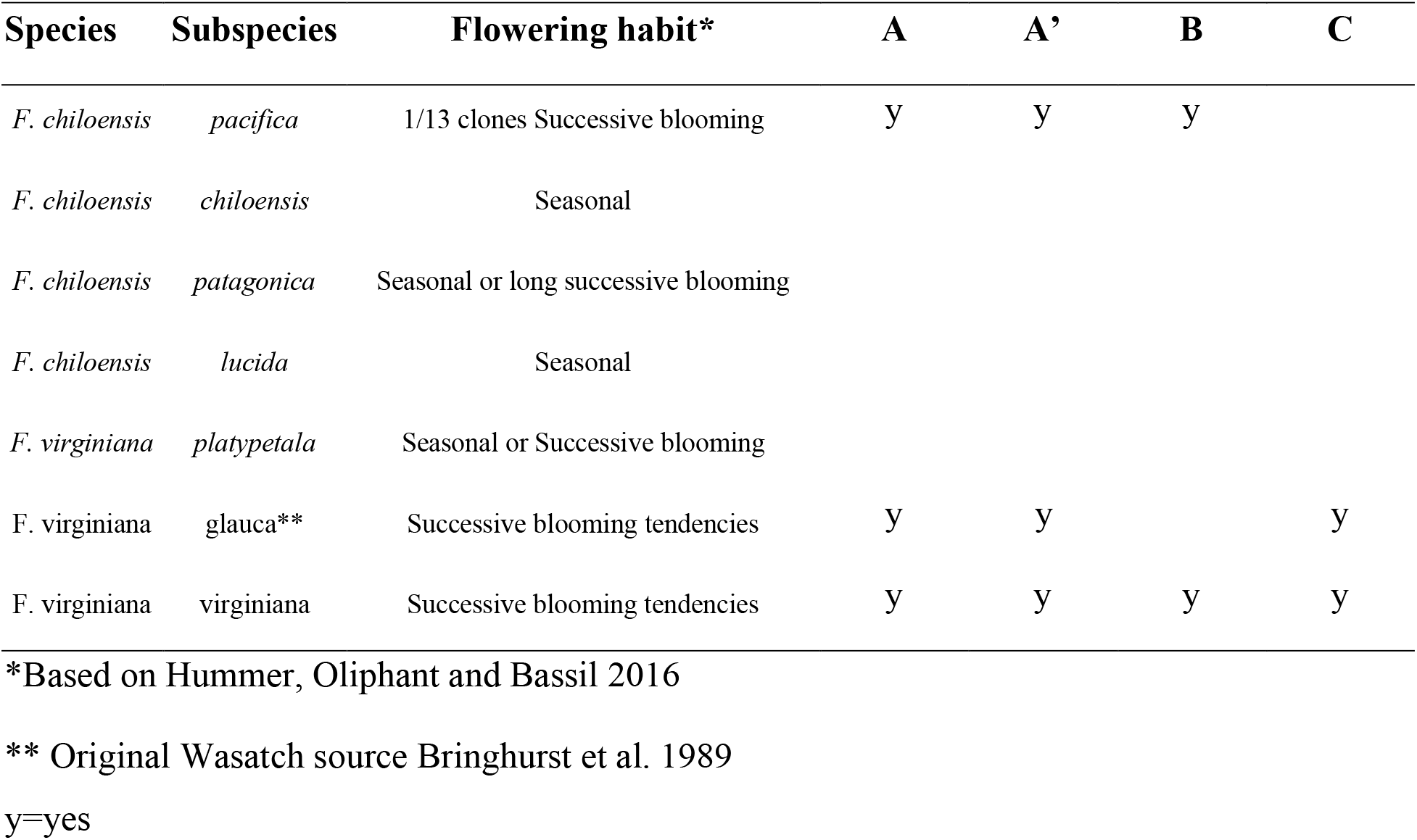
Presence of identified alleles in wild octoploid accessions.

**Figure 5.**
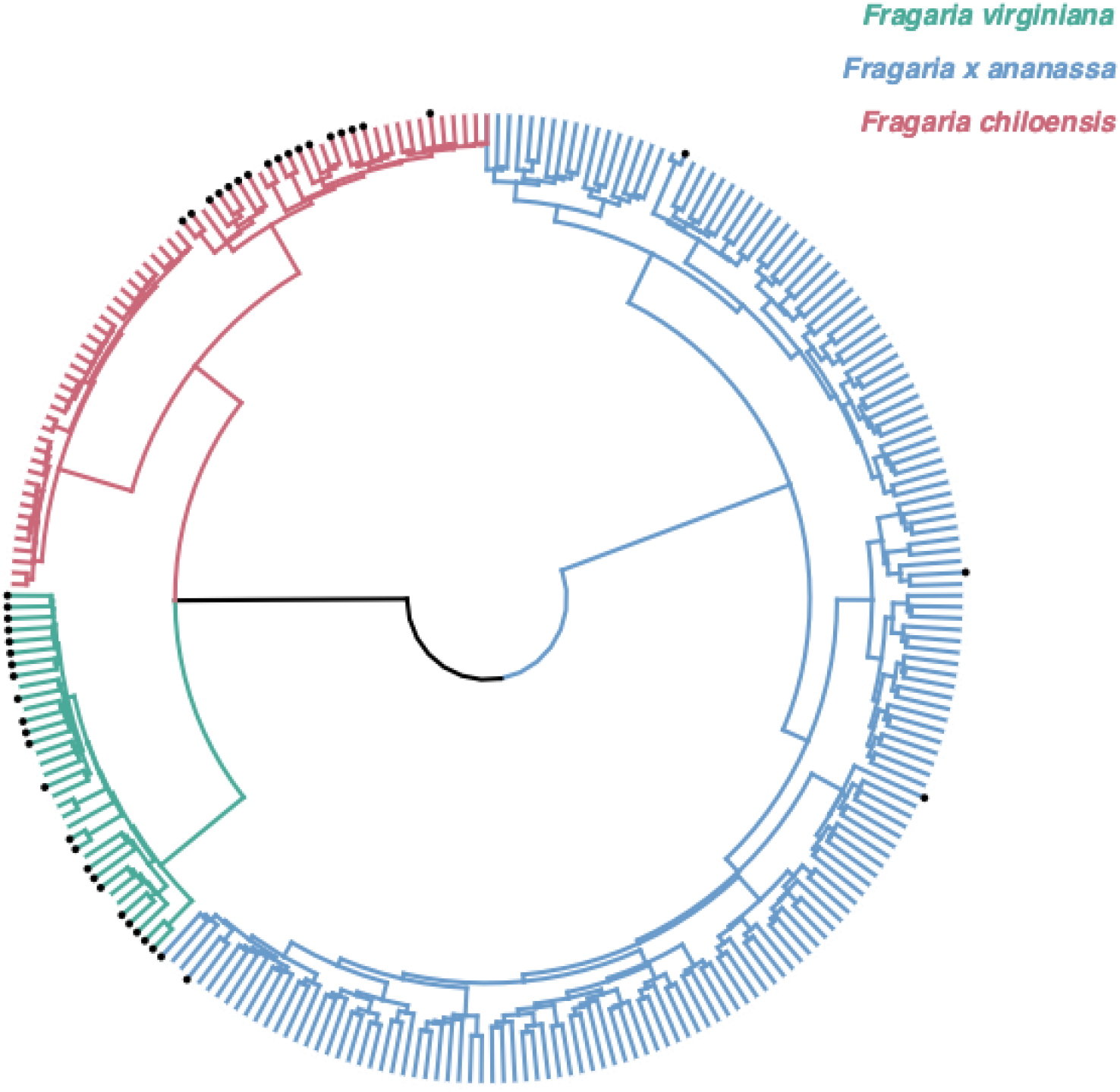
Everbearing trait identified in *Fragaria virginiana* and *Fragaria chiloensis* clades. Dendrogram of *Fragaria* species with everbearing phenotypes highlighted by black nodes; green, *Fragaria virginiana*; pink, *Fragaria chiloensis* and blue, *Fragaria x ananassa.* Data from Hardigan et al (2018).

## Discussion

We show for the first time that complex epistatic interactions across unlinked loci influence expression of the everbearing flowering trait in strawberries. We also provide a model that can be used to increase the efficiency of incorporating the Wasatch source of everbearing into strawberry breeding material.

### The Three Allele Model and Epistatic Interaction Controlling the Everbearing Trait

The major effect QTL, *FaPFRU*, controlling the everbearing trait in cultivated strawberry was previously identified on chromosome 4A (Gaston *et al*., 2013; Perrotte *et al*., 2016a) and has been utilised extensively by breeding companies. It was believed that a single dominant *FaPFRU* allele was sufficient to achieve the everbearing trait, however, spurious segregation patterns have been observed in breeding populations (Lewers *et al*., 2019). Lewers et al. (2019) predicted the hypothetical existence of epistatic enhancer and repressor modulators involved in the expression of the everbearing trait. Our GWAS has revealed three loci associated with the everbearing trait including a major SNP on chromosome 4A, here named A and two potential epistatic enhancers in chromosomes 5C and 6A named B and C, respectively.

The major locus A, on chromosome 4A, is located close to a previously identified region, *FaPFRU* (Hardigan et al., 2018). The previously identified significant SNP named A’ in our study was 3.5 Mb away from A on chromosome 4A and suggested that recombination is strongly suppressed between these two loci in everbearing genotypes, but not in June-bearers. However, an additional SNP called X, located between A and A’, showed a stronger association with the everbearing trait than A’ in our germplasm. Considering the highly significant interaction between the two loci A and X found in this study, it is likely that *FaPFRU* is located between these markers on chromosome 4A. However, the lack of SNP markers in the area prevents a more accurate detection of the locus in our dataset. Previously, *FaPFRU* was mapped between the markers bx089 and bx215 that delimited a QTL region of ~1.1 Mb, corresponding to the genomic fragment between 29,738,228 - 30,924,902 bp in *F. vesca* chromosome 4 (Perrotte et al., 2016a), located just 200 kb downstream of the marker A. Nonetheless, here we have identified eight genotypes with putative recombinations in the region that are potentially useful for the identification of causative alleles.

All individuals containing two copies of the major allele “AA” were everbearers. Also the majority of individuals (98%) that were heterozygotes for the major allele “Aa” but contained two copies of “B” or “C” modifier alleles (either BB or CC, or heterozygous for both loci) were everbearers. The majority of individuals (88%) that did not contain a modifier were June-bearers and individuals containing a single modifier segregated for flowering habit. Based on these findings, we suggest that B and C are functionally redundant quantitative enhancers of *FaPFRU* that are involved in the expression of the everbearing trait, at least in individuals that are heterozygous for dominant *FaPFRU* alleles. Furthermore, the segregation of June-bearing and everbearing phenotypes in the genotypes “AaBbcc'' and “AabbCc” led us to hypothesise that there could be a fourth undetected QTL “D”, but further work is needed to test this hypothesis.

Through predicting the segregation ratios of progeny in our validation crosses, the complex three gene model with epistatic interactions showed the best fit between expected and observed phenotypes. This supports the importance of the two epistatic modifiers identified here. Nonetheless, the *R^2^* value of the model was 0.33 indicating that other loci are likely to be involved in the expression of the everbearing trait, at least within the studied germplasm.

The 18 everbearers that did not contain a copy of the major allele A may have undergone a recombination between the focal maker Affx-88857614 and the causative allele. Alternatively these individuals may contain other “non-Wasatch” sources of the everbearing trait. Indeed, other sources of the everbearing trait have been introgressed into breeding germplasm and described. For example, a QTL on linkage group 3C was identified to influence the intensity of late season flowering (Perrotte *et al., 2016b)*, and a significant locus for everbearing trait was identified on chromosome 3 in the pre-1990 breeding material that lacked the dominant *FaPFRU* locus (Hardigan et al., 2018). It is possible that some of the everbearing but homozygous recessive individuals “aa” identified in this study may contain these sources of continual flowering.

### Molecular Control of Perpetual Flowering

Previous studies have identified candidate genes for *FaPFRU* locus including *FLOWERING LOCUS T2* (FaFT2) and PHYTOCHROME B ACTIVATION TAGGED SUPPRESSOR 1 (BAS1/CYP72B1) (Perrotte et al., 2016a; Hardigan et al., 2018). The role that *FaPFRU* and *BAS1/CYP72B1* play in the control of the everbearing trait remains to be shown, although recent work has provided evidence that FT2 is a florigen with a role as a mobile flower inducing signal (Gaston et al., 2021). The mechanism controlling perpetual flowering has been extensively characterised in the diploid woodland strawberry (*F. vesca* L.) and many interacting components have been described (Koskela *et al.*, 2016). None of these genes are located close to the *FaPFRU* loci, nor the two other loci identified here, but we have found a gene encoding a Phytochrome A photoreceptor less than 100 kb away from the most significant marker on chromosome 6A in *F. vesca* genome.

In *F. vesca*, a mutation in the gene encoding the floral repressor TERMINAL FLOWER1 (TFL1) leads to a perpetual flowering phenotype (Koskela *et al*., 2012), demonstrating that FvTFL1 functions as a floral repressor in this species. Because *FaTFL1* is not located close to an everbearing QTL in cultivated strawberries, we hypothesise that regulators of the everbearing trait either function as suppressors of *FaTFL1* or bypass its repressor function. Three other genes, *CONSTANS* (*FvCO*), another putative florigen gene *FT1* and *SUPPRESSOR OF THE OVEREXPRESSION OF CONSTANS1* (*SOC1*) control photoperiodic flowering in *F. vesca* (Koskela *et al*., 2012, 2016; Rantanen *et al*., 2014, 2015). If FaPFRU functioned as a repressor of *FaTFL1*, the long day-activated photoperiodic pathway genes *FaCO*, *FaFT1* and *FaSOC1* could be involved in rapid activation of flowering in long days, and in short days, normal flower induction could take place after gradual downregulation of *FaTFL1* (Koskela et al., 2016). This hypothetical model could explain the relatively weak photoperiodic responses of everbearing cultivars that are also commonly called as day-neutral plants (Hardigan et al., 2018; Hytönen & Kurokura 2020). It is clear that multiple regulators interact to modulate flowering habits in strawberries according to environmental signals (Rantanen et al., 2014; 2015; Koskela et al., 2016), but additional studies are needed to establish how known regulators and unknown loci are intertwined to control everbearing in cultivated strawberries. Moreover, studies should explore the function of the modifiers in different strawberry production areas, as previous studies have shown that expression of the everbearing trait is influenced by the local environment (Weebadde *et al*., 2007; Castro *et al*., 2015). This knowledge would enable the fine tuning of fruit production in everbearing cultivars within different environments.

### The Origin of the Everbearing Trait

Markers associated with everbearing alleles were found in the wild everbearing octoploid species *F. chiloensis* ssp. *pacifica*, *F. virginiana* ssp. *glauca* and *F. virginiana* ssp. *virginiana*. These findings are consistent with the reported origin of the everbearing phenotype in the studied cultivated strawberry having been sourced from *F. virginiana* ssp. *glauca* (Hancock & Simpson, 1995; Sakin *et al., 1997)*. However, the alleles were also found in a *F. chiloensis* ssp. which could indicate that the everbearing loci arose prior to speciation of *F. chiloensis* and *F. virginiana.* Alternatively, it is possible that gene flow has occurred between the two species and selection has fixed the trait, as *F. chiloensis* ssp. *pacifica* is a northerly subspecies with a similar latitudinal range to *F. virginiana* spp. *glauca*, on the west coast of North America (Hummer *et al., 2011*).

### Trade-off between Flowering and Runner Formation

Vegetative propagation of strawberry cultivars requires the production of runners from axillary buds. However, there is a long standing observation that runner production negatively correlates with flower production (Serçe & Hancock, 2003) This trade-off between flower truss and runner production is particularly clear in everbearing strawberries, because the activation of continuous flowering by FaPFRU causes strong suppression of runner production (Gaston *et al.*, 2013). It has been shown that there is no difference between the number of runners produced by cultivars homozygous or heterozygous for the dominant *FaPFRU* allele (Perrotte *et al.*, 2016b). However, as seen in our work, the phenotype of heterozygous “Aa” is influenced by modifiers B and C, and therefore, their possible effects on runner production should be studied. Ideally, introducing stronger environmental regulation of flowering in everbearing strawberries would allow growers and nurseries to switch between the production of runners and flowers by changing the growing conditions.

## Conclusions

The continuous flowering trait may be partially explained by the three allele model detailed in this study - one large effect allele *FaPFRU* and two modifier alleles, B and C. The modifier alleles are functionally redundant such that two modifiers (either ‘CC’, ‘BB’ or ‘BC’) are required to produce the everbearing flowering habit in individuals containing a single copy of the dominant *FaPFRU* allele. Our results go some way towards explaining why breeders have been observing non-Mendelian segregation in what was once believed to be a monogenic trait.

## Acknowledgements

The authors acknowledge Robert Vickerstaff for generating the octoploid consensus map as part of other projects (Biotechnology and Biological Sciences Research Council (BBSRC) BB/K017071/1; BB/N006682/2 and Innovate UK project 101630) and Dr Beatrice Denoyes, INRA and Dr Amparo Monfort, CRAG for granting the use of their informative markers in the production of the strawberry consensus linkage map. The authors acknowledge funding from the BBSRC BB/M01200X/2, BB/N006682/1 and Innovate UK project 101914. In addition, HC’s time, funded by Growing Kent and Medway (107139).

## Author contributions

HMC, CN, TH, MS, RJH – Conceived and designed experiments

MS, HMC, RJH – Conducted quantitative genetics analysis and marker analysis

CN, TH, KH, AW – Performed experiments

SL – Performed DNA extraction for genotyping

HMC, CN, TH, & RJH – wrote the manuscript with contributions from all authors.

## Data Availability Statement

Data will be made available upon reasonable request through contacting the corresponding author.

---

**Supplementary Figure 1.**
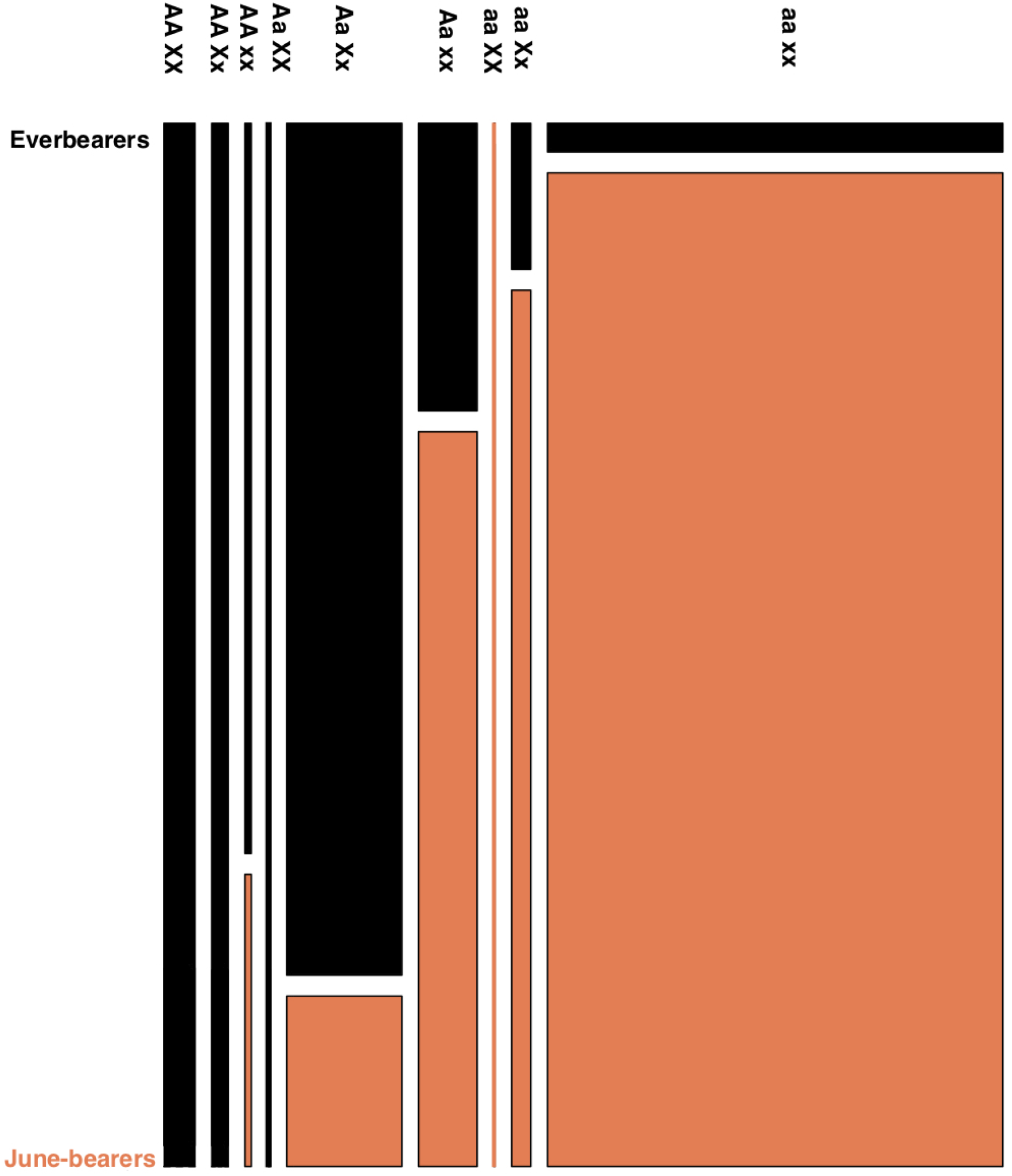
Phenotypic proportion of individuals for each genotype across the gap at the distal portion of Chromosome 4A displayed as a mosaic plot. The width of the block is proportional to the number of individuals with a given genotype. ‘A’ represents the major allele associated with everbearing (Affx-88857614) ‘X’ represents the most significant marker associated with everbearing after the 2.85 Mb gap (Affx-88857910). Pink (June-bearers) and black (Everbearers) box sizes represent the proportion of phenotypes in each genotype class.

## References

Andrés F, Coupland G. 2012. The genetic basis of flowering responses to seasonal cues. Nature Reviews. Genetics 13: 627–639.

Bassil NV, Davis TM, Zhang H, Ficklin S, Mittmann M, Webster T, Mahoney L, Wood D, Alperin ES, Rosyara UR, et al. 2015. Development and preliminary evaluation of a 90 K Axiom® SNP array for the allo-octoploid cultivated strawberry Fragaria × ananassa. BMC Genomics 16: 1310.

Bouché F, Woods DP, Amasino RM. 2017. Winter Memory throughout the Plant Kingdom: Different Paths to Flowering. Plant Physiology 173: 27–35.

Bradford E, Hancock JF, Warner RM. 2010. Interactions of temperature and photoperiod determine expression of repeat flowering in strawberry. Journal of the American Society for Horticultural Science 135: 102–107.

Bringhurst R, Voth V. 1980. Six new strawberry varieties released. California agriculture.

Bringhurst RS, Voth V. 1984. Breeding octoploid strawberries. Iowa State J. Res.

Castro P, Bushakra JM, Stewart P, Weebadde CK, Wang D, Hancock JF, Finn CE, Luby JJ, Lewers KS. 2015. Genetic mapping of day-neutrality in cultivated strawberry. Molecular Breeding 35: 79.

Darrow GM. 1966. The strawberry. History, breeding and physiology. The strawberry. History.

Faedi W, Mourgues F, Rosati C. 2002. Strawberry breeding and varieties: situation and perspectives. Acta horticulturae: 51–59.

Galili T. 2015. dendextend: an R package for visualizing, adjusting and comparing trees of hierarchical clustering. Bioinformatics 31: 3718–3720.

Gaston A, Perrotte J, Lerceteau-Köhler E, Rousseau-Gueutin M, Petit A, Hernould M, Rothan C, Denoyes B. 2013. PFRU, a single dominant locus regulates the balance between sexual and asexual plant reproduction in cultivated strawberry. Journal of Experimental Botany 64: 1837–1848.

Gaudinier A, Blackman BK. 2020. Evolutionary processes from the perspective of flowering time diversity. The New Phytologist 225: 1883–1898.

Guttridge CG. 1985. Fragaria ananassa. CRC handbook of flowering.

Hancock J, Simpson D. 1995. Methods of extending the strawberry season in europe. HortTechnology 5: 286–290.

Hardigan MA, Poorten TJ, Acharya CB, Cole GS, Hummer KE, Bassil N, Edger PP, Knapp SJ. 2018. Domestication of Temperate and Coastal Hybrids with Distinct Ancestral Gene Selection in Octoploid Strawberry. The plant genome 11.

Hummer KE, Bassil N, Njuguna W. 2011. Fragaria. In: Kole C, ed. Wild crop relatives: genomic and breeding resources. Berlin, Heidelberg: Springer Berlin Heidelberg, 17–44.

Koskela EA, Mouhu K, Albani MC, Kurokura T, Rantanen M, Sargent DJ, Battey NH, Coupland G, Elomaa P, Hytönen T. 2012. Mutation in TERMINAL FLOWER1 reverses the photoperiodic requirement for flowering in the wild strawberry Fragaria vesca. Plant Physiology 159: 1043–1054.

Koskela EA, Sønsteby A, Flachowsky H, Heide OM, Hanke M-V, Elomaa P, Hytönen T. 2016. TERMINAL FLOWER1 is a breeding target for a novel everbearing trait and tailored flowering responses in cultivated strawberry (Fragaria × ananassa Duch.). Plant Biotechnology Journal 14: 1852–1861.

Krizek BA, Fletcher JC. 2005. Molecular mechanisms of flower development: an armchair guide. Nature Reviews. Genetics 6: 688–698.

Lewers KS, Castro P, Hancock JF, Weebadde CK, Die JV, Rowland LJ. 2019. Evidence of epistatic suppression of repeat fruiting in cultivated strawberry. BMC Plant Biology 19: 386.

Liston A, Cronn R, Ashman T-L. 2014. Fragaria: a genus with deep historical roots and ripe for evolutionary and ecological insights. American Journal of Botany 101: 1686–1699.

Mookerjee S, Mathey MM, Finn CE, Zhang Z, Hancock JF. 2013. Heat tolerance plays an important role in regulating remontant flowering in an F1 population of octoploid strawberry (Fragaria×ananassa). Journal of berry research 3: 151–158.

Nellist, CF, Vickerstaff RJ, Sobczyk MK, Marina-Montes C, Wilson FM, Simpson DW, Whitehouse AB, Harrison RJ. 2019. Quantitative trait loci controlling *Phytophthora cactorum* resistance in the cultivated octoploid strawberry (*Fragaria* x *ananassa*). Horticultural Research 6: 1–14.

